# hnRNPA1 Remodels SARS-CoV-2 Genomic G-Quadruplexes through Its UP1 Domain

**DOI:** 10.64898/2026.10.16.759555

**Authors:** Parul Gupta, Swarnava Majumdar, Aditi Maulik, Mahavir Singh

## Abstract

The single-stranded RNA genome of SARS-CoV-2 contains several predicted G-quadruplex (G4) forming sequences. Pharmacological stabilisation of these structures has been shown to suppress viral replication, suggesting that their resolution may be required for efficient viral replication. However, the roles of host proteins in stabilization or destabilization of G4 structures have remained largely unexplored. Human hnRNPA1, one of the most abundant RNA-binding proteins in eukaryotic cells, is a G-quadruplex unfolding protein and has independently been identified as a SARS-CoV-2 genome-binding protein. However, whether it acts on the G-quadruplexes within the SARS-CoV-2 genome has not been examined. Here, we screened eight putative G-quadruplex forming sequences conserved across SARS-related coronaviruses and identified two sequences that fold into G4 structures, confirmed by circular dichroism (CD) spectroscopy and thioflavin T (ThT) fluorescence-based experiments. Furthermore, we showed that the UP1 and UP1+RGG domains of hnRNPA1 specifically bind and unfold these two viral RNA G-quadruplexes. Both UP1 and UP1+RGG unfold viral RNA G-quadruplexes to similar extent, indicating that recognition and unfolding of these viral G-quadruplexes are mediated mainly by the UP1 domain of hnRNPA1. Together, these findings identify hnRNPA1 as a host factor capable of binding and remodelling SARS-CoV-2 genomic RNA G-quadruplexes and suggest a mechanism by which the virus may exploit an abundant host protein to resolve G4 structures within its genome.

## Introduction

G-quadruplexes (G4s) are one of the non-canonical secondary structures that are formed by the guanine-rich regions of DNA or RNA [1]. They consist of two or more stacked G-tetrads, each comprising four guanines connected by Hoogsteen hydrogen bonds and stabilised by monovalent cations such as K^+^or Na^+^[1,2]. G4s have been visualised in both the nucleus and cytoplasm of human cells, where they regulate processes including telomere maintenance, DNA replication, transcription, alternative splicing, and translation [3–12].

G4s have emerged as potential drug targets against pathogenic viruses because of their occurrence in various viral genomes and their regulatory roles in the viral life cycle [13]. Previous studies have shown roles of G4s in human viruses such as human immunodeficiency virus (HIV), herpes simplex virus (HSV), Epstein-Barr virus (EBV), human papillomavirus (HPV), hepatitis B virus (HBV), Nipah virus, hepatitis C virus (HCV), Zika virus, Ebola virus and SARS-CoV-2 [13–19]. In SARS-CoV-2, multiple potential G4-forming sequences have been predicted on both positive- and negative-sense RNA strands [20,21,30,31,22–29], and several have been experimentally validated *in vitro* [17,24,26,30,31]. G4-stabilising ligands have been shown to inhibit SARS-CoV-2 infection by targeting viral RNA G4s and by stabilising G4 within the host TMPRSS2 transcript, supporting G4s as potential therapeutic targets [17,32].

Several viral and host proteins have been reported to interact with SARS-CoV-2 RNA G4s. The viral helicase nsp13 was shown to bind SARS-CoV-2 G4s and distort their structure [24], whereas the SARS-unique domain (SUD) of nsp3 was shown to recognise and stablize RNA G4s *in vitro* [33]. Host factors including CNBP, RHAU53 peptide of DHX36, and human DNA topoisomerase I have also been shown to interact with viral G4s, highlighting the potential importance of host proteins in regulating their structure and accessibility [25,28,34].

Human hnRNPA1 is a highly abundant nucleocytoplasmic shuttling protein [35,36]. It comprises a folded N-terminal UP1 (unwinding protein 1) domain containing two RNA recognition motifs (namely RRM1 and RRM2), followed by a C-terminal intrinsically disordered low-complexity domain enriched in glycine and arginine residues containing RGG-box, a prion-like domain and a M9 shuttling sequence that directs its bidirectional nucleocytoplasmic transport [37,38]. hnRNPA1 is a multifunctional RNA-binding protein involved in diverse aspects of RNA metabolism, including transcriptional regulation, alternative pre-mRNA splicing, mRNA transport and localisation, stability, translation, and miRNA processing [36,39–44].

hnRNPA1 preferentially binds single-stranded, guanine-rich sequences and destabilises the higher-order G4 structures mainly using its UP1 and RGG-box domains [9,37,45]. hnRNPA1 unfolds the G4 by binding the unfolded state with higher affinity and thereby drawing the folding equilibrium away from the folded G-quadruplex structure [37]. hnRNPA1 has been shown to act on telomeric DNA and TERRA RNA G-quadruplexes as well as G-quadruplexes in the promoter regions of KRAS and *TRA2B*, and in the 5′ UTRs of *RON, FEN1* and *KRAS*, influencing telomere maintenance, transcription, translation and mRNA stability [39,40,43,44,46]. The N-terminal UP1 domain carries the G4 unfolding activity and has been shown to destabilise both DNA and RNA G-quadruplexes, although the efficiency of unfolding depends on the topology of the substrate [9,37,45]. The central RGG-box was shown to bind folded G-quadruplexes but not single-stranded nucleic acid and enhances UP1-mediated unfolding of telomeric DNA and RNA G-quadruplexes [9,45].

hnRNPA1 has also been implicated in the regulation of viral RNA G4s. In Kaposi’s sarcoma-associated herpesvirus (KSHV), hnRNPA1 was shown to unfold a G4 within LANA mRNA and promotes translation [47]. In PRRSV-2, the UP1 domain was shown to mediate recognition of a guanine-rich sequence in antisense RNA [48]. hnRNPA1 was also shown to bind coronavirus RNAs such as in mouse hepatitis virus (MHV), it associates with transcription-regulatory and 3′ untranslated regions and modulates viral RNA synthesis [49,50]. Genome-wide interactome studies further identified hnRNPA1 as an interactor of SARS-CoV-2 RNA, and hnRNPA1 has been implicated in regulating the transition from early translation to replication of the viral genome through recognition of m6A-modified RNA [51,52].

Despite increasing evidence for the existence and functional significance of SARS-CoV-2 RNA G4s, the mechanisms by which host RNA-binding proteins recognise and remodel these structures remain poorly understood. In particular, it is unclear whether hnRNPA1 directly recognises G4-forming regions within the SARS-CoV-2 genome and how such interactions influence G4 structure. In this study, we investigated the interaction and unfolding of putative SARS-CoV-2 RNA G-quadruplexes by hnRNPA1. Conserved putative G4-forming sequences were identified in the Wuhan reference genome and screened for G4 formation. Two sequences, located at genomic positions 3467 and 13385 were experimentally validated to form G4s structures. Isothermal titration calorimetry showed that both the UP1 and UP1+RGG domains of hnRNPA1 bind these viral G4s with low-micromolar affinity. Circular dichroism spectroscopy and thioflavin T fluorescence assays demonstrated that both constructs efficiently unfold the viral G4s to a similar extent, indicating that the UP1 domain is the principal determinant of G4 recognition and unfolding. Together, our results identify hnRNPA1 as a host factor capable of remodelling SARS-CoV-2 RNA G-quadruplexes and suggest that this activity is mediated predominantly by its UP1 domain.

## Materials and Methods

### Prediction of G-quadruplexes in SARS-CoV-2 genome

Complete genome sequences of four different strains of SARS-CoV-2 (Wuhan, Italy, USA, and India) were aligned in the Clustal Omega multiple sequence alignment program [53]. They were found to be >99% conserved. Wuhan sequence (NCBI reference number: NC_045512.2 and GI 1798174254) was used for mapping putative G-quadruplexes in QGRS Mapper. QGRS Mapper is a software program that generates information on the composition and distribution of putative Quadruplex forming G-rich Sequences (QGRS) in nucleotide sequences [54]. It returned 25 non-overlapping potential G4-forming sequences in the complete genome of SARS-CoV-2. Next, a comprehensive database of G-quadruplex forming sequences in the genome of all known human viruses was searched for the SARS group of viruses. The list of all G4s present in the SARS genome, their positions, and the degree of conservation of each G4s in its genome context was retrieved for the SARS group of viruses. Extensive analysis of the database revealed 11 conserved G4 sequences as characteristics of the coronavirus (-ve RNA strand) genome. Multiple sequence analysis of SARS coronavirus sequence (NCBI Genebank number GU553365.1) with the SARS-CoV-2 sequences identified that out of the 11 conserved characteristic G4s in the SARS coronavirus dataset, eight sequences were found to be absolutely conserved in SARS-CoV-2 strains. Then the G4s were mapped with respect to the different protein-coding parts of the virus. Several of the predicted G4 sequences were found to be in the coding region of viral proteins.

### DNA and RNA oligomer sequences

The following table contains the nucleic acid sequences used in this study. The DNA and RNA sequences were ordered from Sigma-Aldrich.

### RNA G-Quadruplex stabilising compounds

The ligands used for stabilising the G-quadruplex topology were BRACO-19 hydrochloride and TMPyP4 tosylate (5,10,15,20-tetrakis(1-methyl-4-pyridyl)porphyrin tetra(p-toluenesulfonate)). Both compounds were purchased from Sigma-Aldrich.

### G-quadruplex preparation

A stock concentration of 1 mM DNA/RNA was prepared by dissolving the lyophilised oligos in nuclease-free water. Working concentrations were prepared by diluting the stock in the appropriate buffer (20 mM HEPES pH 7.4, 100 mM KCl) for subsequent experiments.

To form the G-quadruplex, samples were heated at 95 °C for 4 minutes and allowed to cool gradually at room temperature for 2 hours. The prepared G-quadruplex stocks were stored at −20 °C until further use. The concentration of oligos was determined using a UV spectrophotometer at 260 nm, and molar extinction coefficients were calculated based on the sequence of the respective oligos.

### Protein expression and purification

hnRNPA1 (UniProt identifier P09651-2) protein constructs, UP1 (residue 1-196) and UP1+RGG (residue 1-249), were expressed in BL21 (DE3) Rosetta cells grown in LB Medium as per the established protocols [9]. Briefly, the primary culture was grown overnight at 37 °C and was used to inoculate the secondary culture. After inoculation, the culture was allowed to grow until the OD_600_ nm reached ∼0.6. The culture was induced with 0.5 mM IPTG and incubated overnight at 30 °C. Cells were harvested by centrifuging at 4500 rpm for 15 minutes at 4 °C. All the subsequent purifications were performed at 4 °C. The cell pellet was resuspended in lysis buffer (25 mM Tris, pH 7.4, 500 mM NaCl, 5 mM MgCl_2_, 10 % Glycerol and 5 mM β-mercaptoethanol) and lysed by sonication. The lysate was then centrifuged at 13000 rpm for 60 minutes. The supernatant was collected and loaded onto a Ni^2+^-NTA column for affinity-based purification. The column was first washed with lysis buffer containing 5 mM imidazole, then with lysis buffer supplemented with 20 mM imidazole. The elution of the protein of interest was conducted using an elution buffer (lysis buffer + 400 mM Imidazole). Eluted fractions were pooled and subjected to two rounds of dialysis, each lasting 6 hours. Following dialysis, the protein was further purified using cation-exchange chromatography with buffer A (25 mM Tris, pH 8.0, 50 mM NaCl) and buffer B (25 mM Tris, pH 8.0, 1 M NaCl). Fractions containing the protein were collected, concentrated using centrifugal filters, and finally purified by size exclusion chromatography (SEC) on an S75 column equilibrated buffer comprising 20 mM HEPES pH 7.4, with 100 mM KCl.

### Circular Dichroism (CD)

10 μM DNA/RNA samples were prepared in 20 mM HEPES buffer (pH 7.4) and 100 mM KCl, then folded to form a G-quadruplex and subsequently analysed. Changes in the quadruplex structures of DNA and RNA upon interaction with UP1 and UP1+RGG were monitored using CD spectroscopy. These CD studies were carried out on a JASCO J-815 spectrophotometer. The titration experiments were performed in a 1 mm path length cuvette (Hellma Analytics) at 25 °C with a scanning speed of 100 nm/min and a response time of 4 s. Spectra were recorded over a wavelength range of 320 nm to 220 nm. 10 μM of the DNA/RNA was titrated with increasing protein concentrations in molar ratios of 1:0, 1:0.5, 1:1 and 1:2. The average of three spectral scans was taken for each sample. Baseline correction was performed for each sample to subtract the buffer signal. The data were normalised to molar ellipticity per residue. The data obtained were plotted and analysed using OriginPro 8.5. Relative foldedness was calculated by dividing the ellipticity at 264 nm of each protein-containing sample by the corresponding value obtained in the absence of protein and is expressed as a percentage.

### CD thermal melting experiments

To conduct melting experiments on RNA G-quadruplexes with G-quadruplex-stabilising molecules, a CD scan was performed at 5 °C intervals. The scan was initiated at 20 °C and ended at 95 °C. 10 μM samples of RNA G-quadruplexes with the stabilising compounds were incubated in buffer containing 20 mM HEPES (pH 7.4) and 100 mM KCl at room temperature in a 1:1 stoichiometry for 10-15 minutes before conducting the scan. The ellipticity values were taken from the wavelength corresponding to the maximum ellipticity, which was at 264 nm in the spectrum. The melting temperature was calculated by fitting the melting curves to a Boltzmann sigmoidal equation using GraphPad Prism.

### Thioflavin T (ThT) assay

Thioflavin T (ThT; Sigma-Aldrich, T3516) was used for the fluorescence-based detection of G-quadruplex formation. Fluorescence measurements were carried out in a 96-well microplate to monitor ThT fluorescence emission. For validation of putative G-quadruplex formation, pre-folded G-quadruplex oligonucleotides were incubated with ThT in the buffer 20 mM HEPES, pH 7.4, 100 mM KCl. The final reaction mixture contained 2 μM G-quadruplex oligonucleotide and 1 μM ThT in a total volume of 200 μl per well and was incubated for 20 minutes before measuring the emission. To examine G-quadruplex unfolding by UP1 and UP1+RGG, proteins were added to the pre-folded G-quadruplex samples at a working concentration of 2 μM, while maintaining the oligonucleotide and ThT concentrations constant as described above. All experiments were performed in triplicate.

Fluorescence spectra were recorded with an excitation wavelength of 425 nm, and emission was monitored over the range of 450–700 nm. All measurements were performed in three independent experiments, and fluorescence values were corrected against a buffer-only control containing ThT. The data obtained were plotted and analysed using GraphPad Prism. Relative foldedness was calculated by dividing the ThT fluorescence intensity at 485 nm of each protein-containing sample by the corresponding value obtained in the absence of protein, and is expressed as a percentage.

### Isothermal titration calorimetry (ITC)

ITC experiments were performed using a Microcal PEAQ-ITC instrument at 25 °C. All the samples and buffers were thoroughly degassed before running the experiment. Both protein and RNA samples were prepared in a buffer containing 20 mM HEPES, pH 7.4, and 100 mM KCl. For titration, the sample cell was filled with 10-30 μM RNA and the syringe was filled with 100-300 μM protein. A total of 19 injections of titrant were performed at an interval of 180 s between successive injections. The heat of dilution was subtracted from the raw integrated heat data, and the resulting data were fit using a one-site binding model. The data were processed and analysed using the software provided by the manufacturer.

## Results and discussion

### Bioinformatics identification and characterisation of putative G-quadruplex forming sequences from the genome of SARS-CoV-2

Our bioinformatic analysis showed presence of 25 putative G-quadruplex forming sequences in the SARS-CoV-2 genome (Supplementary Figure S1). Using different algorithms and rule sets, several previous studies had also predicted some of these putative G4 forming sequences in SARS-CoV-2 RNA [20,21,24,30]. Putative G-quadruplexes have been mapped in both the positive and negative strands and across all seven human coronaviruses [26–28]. Of the 25 predicted sequences, eight sequences were found to be absolutely conserved across SARS-CoV-2 strains and the SARS group of viruses. Therefore, we selected these eight sequences for their further characterisation and functional analysis in this study (Table 1). To validate the ability of the predicted sequences to form G-quadruplex structures *in vitro*, we employed circular dichroism (CD) spectroscopy and Thioflavin T (ThT) fluorescence assays. Different topologies of G-quadruplexes show distinct peaks at particular wavelengths in the CD spectra. Generally, parallel G-quadruplexes exhibit a positive peak at ∼264 nm and a negative peak at ∼245 nm, antiparallel structures show a positive peak at ∼295 nm and a negative peak at ∼265 nm, while hybrid topologies display two positive maxima at ∼295 nm and ∼264 nm along with a negative peak at ∼245 nm [55].

**Table 1.**
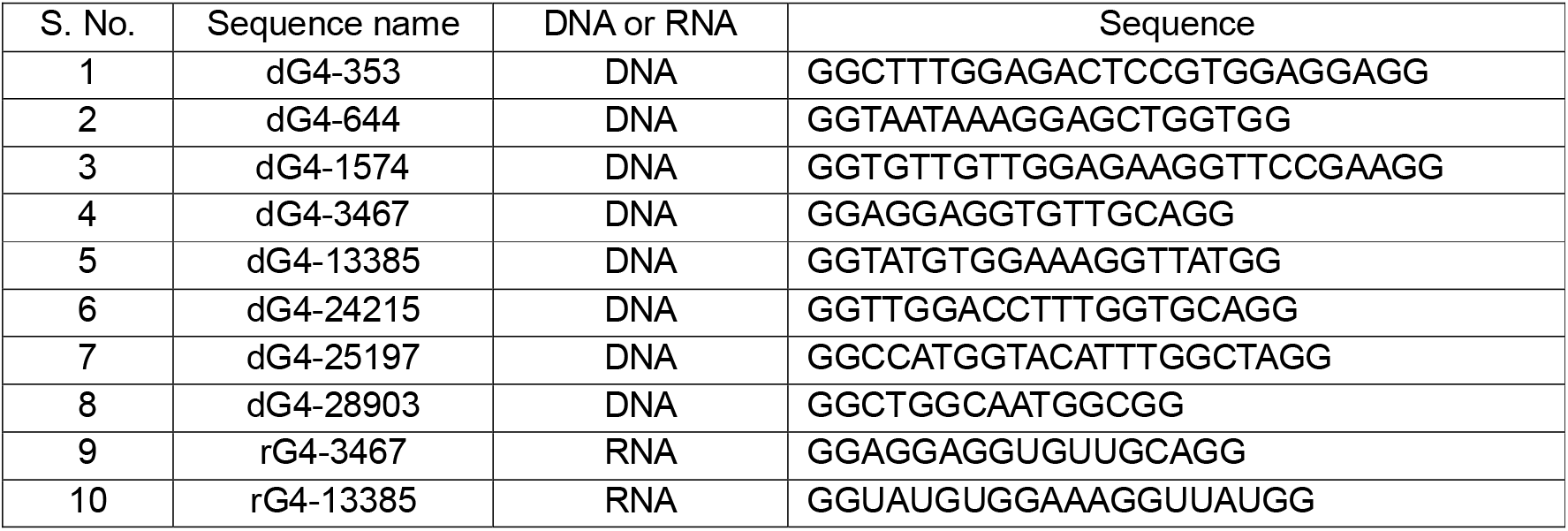
G-quadruplex DNA and RNA oligonucleotides used in this study.

We initiated our studies with all eight putative and conserved sequences first using DNA oligonucleotides. Our CD analysis showed that only two out of the eight sequences, namely 3467 (named dG4-3467) and 13385 (named dG4-13385), exhibited characteristic signatures of G-quadruplex formation (Figure 1A and Supplementary Figure S2). Under 100 mM KCl in the buffer, dG4-3467 showed a positive maximum at ∼264 nm and a negative minimum near ∼240 nm, consistent with a parallel G-quadruplex topology. In contrast, under same conditions, dG4-13385 displayed two positive ellipticity maxima at ∼295 and ∼240 nm, with a negative minimum at ∼264 nm, which is characteristic of an antiparallel G-quadruplex topology. Further to confirm whether the spectra of the individual sequences alter by K^+^ ions, we have conducted CD studies of all eight DNA sequences in the presence and absence of KCl (Supplementary Figure S2). The results showed that only dG4-3467 and dG4-13385 showed enhancement in the ellipticity in CD spectra in the presence of K^+^ ions, again confirming that these two sequences form G-quadruplex structures.

**Figure 1.**
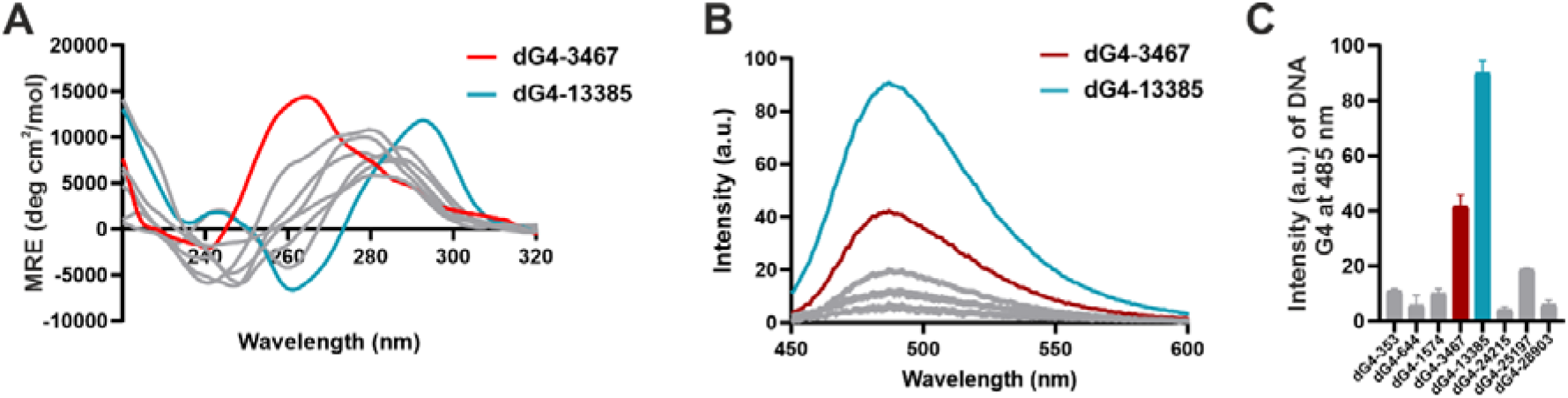
*In vitro* validation of G-quadruplex formation by the eight conserved sequences as DNA oligonucleotides. **(A)** CD spectra of the eight putative DNA G4 oligonucleotides recorded from 220 to 320 nm. dG4-3467 (red) exhibits a positive maximum at ∼264 nm and a negative minimum at ∼240 nm, consistent with a parallel G-quadruplex topology, whereas dG4-13385 (cyan) exhibits positive maxima at ∼295 nm and ∼240 nm with a negative minimum at ∼264 nm, characteristic of an antiparallel G-quadruplex. The remaining six DNA sequences (grey) show no spectral features indicative of quadruplex formation. Spectra are plotted as mean residue ellipticity (MRE) and represent the average of three scans, baseline-corrected against buffer. **(B)** ThT fluorescence emission spectra of the same eight oligonucleotides recorded from 450 to 600 nm following excitation at 425 nm. Consistent with the CD data, dG4-13385 (cyan) and dG4-3467 (red) give higher emission at ∼485 nm than the remaining six sequences (grey). **(C)** ThT emission intensity at 485 nm for each of the eight oligonucleotides, plotted as a bar graph with dG4-3467 (red) and dG4-13385 (cyan) highlighted. Values were corrected against a buffer-plus-ThT control. Data represent the mean of three independent measurements; error bars denote standard deviation with n = 3.

Next, we employed the ThT fluorescence assay to confirm the results obtained from the CD spectroscopy. Thioflavin T (ThT) is a benzothiazole dye widely used as a fluorescent probe to detect G-quadruplex DNA/RNA structures. It binds G-quadruplex through end-stacking, resulting in the restriction of the dye’s internal rotation, leading to an increase in fluorescence intensity that signals the presence and folding of the G-quadruplex [56,57]. The results of the ThT assays are shown in Figure 1B, where the emission spectra of all eight sequences are plotted over the range 450-600 nm. Consistent with the CD studies, in the presence of ThT, the intensities of emission peak of dG4-13385 and dG4-3467 were higher than those of the other DNA sequences at a wavelength of 485 nm (Figure 1B and 1C). This again confirms that these two sequences form G-quadruplex structures.

### SARS-CoV-2 RNA G-quadruplexes adopt parallel topology

As the DNA quadruplexes dG4-3467 and dG4-13385 showed G-quadruplex formation, their corresponding RNA counterparts (rG4-13385 and rG4-3467) were used for further analysis and experiments (Table 1). Both RNA G-quadruplexes exhibited a characteristic CD signal corresponding to a parallel G-quadruplex topology as shown in Figure 2. In the presence of increasing KCl concentration (0, 100, 150 mM), enhancement in the intensity of ellipticity at 264 nm was observed for both rG4-13385 and rG4-3467, clearly showing a K^+^-dependent G-quadruplex formation by these two sequences. Upon addition of KCl, the spectrum shifted to the characteristic spectrum of a parallel G-quadruplex, with increased intensity at 264 nm for both the sequences (Figure 2A and 2B).

**Figure 2.**
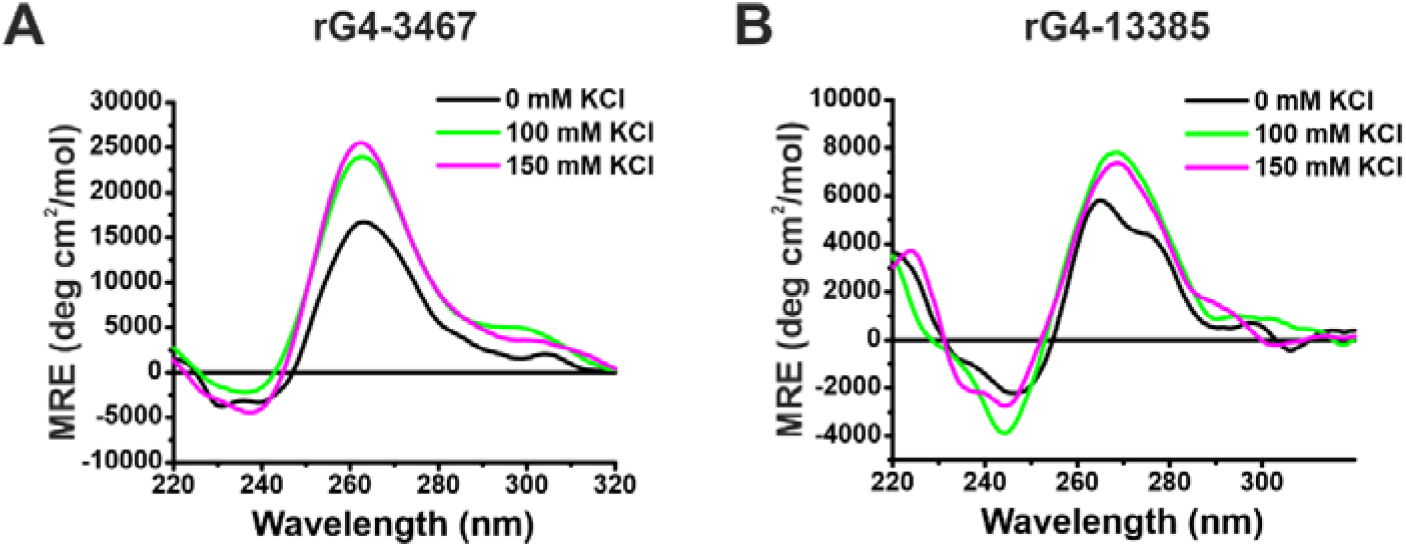
The RNA G-quadruplex-forming sequences adopt a parallel topology and are stabilised by KCl. **(A and B)** CD spectra of rG4-3467 (A) and rG4-13385 (B) recorded from 220 to 320 nm at 0 mM (black), 100 mM (green) and 150 mM (magenta) KCl. In the presence of KCl, the spectrum shows a positive maximum at ∼264 nm and a negative minimum at ∼240 nm, characteristic of a parallel G-quadruplex topology. Spectra are plotted as mean residue ellipticity (MRE) and represent the average of three scans, baseline-corrected against buffer.

To further validate the formation of G-quadruplex by rG4-3467 and rG4-13385, we conducted thermal melting experiments of these two RNA oligomers in the presence of G-quadruplex stabilising compounds: BRACO-19 and TMPyP4. BRACO-19 is a cationic 3,6,9-trisubstituted acridine compound, and TMPyP4 is a cationic porphyrin compound. They are widely used G-quadruplex-interactive ligands. Both bind G-quadruplexes principally through π-stacking. BRACO-19 stacks on a terminal G-tetrad of a parallel-folded quadruplex, whereas TMPyP4 binds externally, stacking on exposed loop and flanking bases [58,59]. The experiment was conducted at a 1:1 RNA:compound stoichiometry (Supplementary Figure S3). Upon the addition of BRACO-19 to rG4-3467, the melting temperature increased from 55.7 °C to 74.9 °C. Upon addition of TMPyP4, the melting temperature increased from 55.7 °C to 69.8 °C (Supplementary Figure S3A). In case of rG4-13385, the addition of BRACO-19 increased the melting temperature from 36.6 °C to 42.9 °C, and the addition of TMPyP4 increased from 36.6 °C to 45.9 °C (Supplementary Figure S3B). These results clearly show these G4 stabilizing BRACO-19 and TMPyP4 stabilize rG4-3467 and rG4-13385, further confirming G4 formation by these two sequences.

### UP1 and UP1+RGG domains of hnRNPA1 interact with SARS-CoV-2 genomic G-quadruplexes

The N-terminal UP1 domain of hnRNPA1 is the well-established G4 binding and unfolding domain of hnRNPA1. The central RGG-box was shown to contribute to the specific binding to the telomeric DNA and TERRA RNA G-quadruplexes [9,45]. Therefore, in next sets of experiments, we tested binding of UP1 and UP1+RGG domains to rG4-3467 and rG4-13385 RNA G-quadruplexes, using isothermal titration calorimetry (ITC) experiments.

In all experiments, RNA in the cell was titrated with either UP1 or UP1+RGG proteins present in the syringe. Figure 3 shows the raw and fitted ITC isotherms, and Table 2 summarises the results obtained for each titration. The ITC isotherms show exothermic binding of both rG4-3467 and rG4-13385 to UP1 and UP1+RGG. The obtained dissociation constants suggest that both RNA G-quadruplexes interact with comparable affinities with the UP1 and UP1+RGG domains of hnRNPA1 (Table 2). For rG4-3467, UP1 bound with a K_d_ of 1.36 ± 0.25 μM, and addition of the UP+RGG domain resulted in a similar affinity, with a K_d_ of 1.10 ± 0.25 μM (Table 2 and Figure 3A and 3B). Similarly, rG4-13385 bound UP1 with a K_d_ of 1.61 ± 0.58 μM, whereas the UP1+RGG domain showed a similar K_d_ of 1.14 ± 0.24 μM for this RNA (Table 2 and Figure 3C and 3D).

**Table 2.** Thermodynamic parameters derived from ITC experiments for the titration of UP1 and UP1+RGG into RNA G-quadruplexes rG4-3467 and rG4-13385.

| S. No. | Experiment | $K_d$<br>($\mu\text{M}$) | $\Delta G$<br>(kcal/mol) | $\Delta H$<br>(kcal/mol) | $-T\Delta S$<br>(kcal/mol) | N |
| --- | --- | --- | --- | --- | --- | --- |
| 1 | UP1:rG4-3467 | $1.36 \pm 0.25$ | -8.01 | $-23.3 \pm 1.08$ | 15.3 | 1.3 |
| 2 | UP1+RGG:rG4-3467 | $1.10 \pm 0.25$ | -8.13 | $-12.8 \pm 0.52$ | 4.69 | 1 |
| 3 | UP1:rG4-13385 | $1.61 \pm 0.58$ | -7.9 | $-26.4 \pm 2.73$ | 18.5 | 1.1 |
| 4 | UP1+RGG:rG4-13385 | $1.14 \pm 0.24$ | -8.11 | $-38.5 \pm 2.24$ | 30.4 | 0.817 |

**Figure 3.**
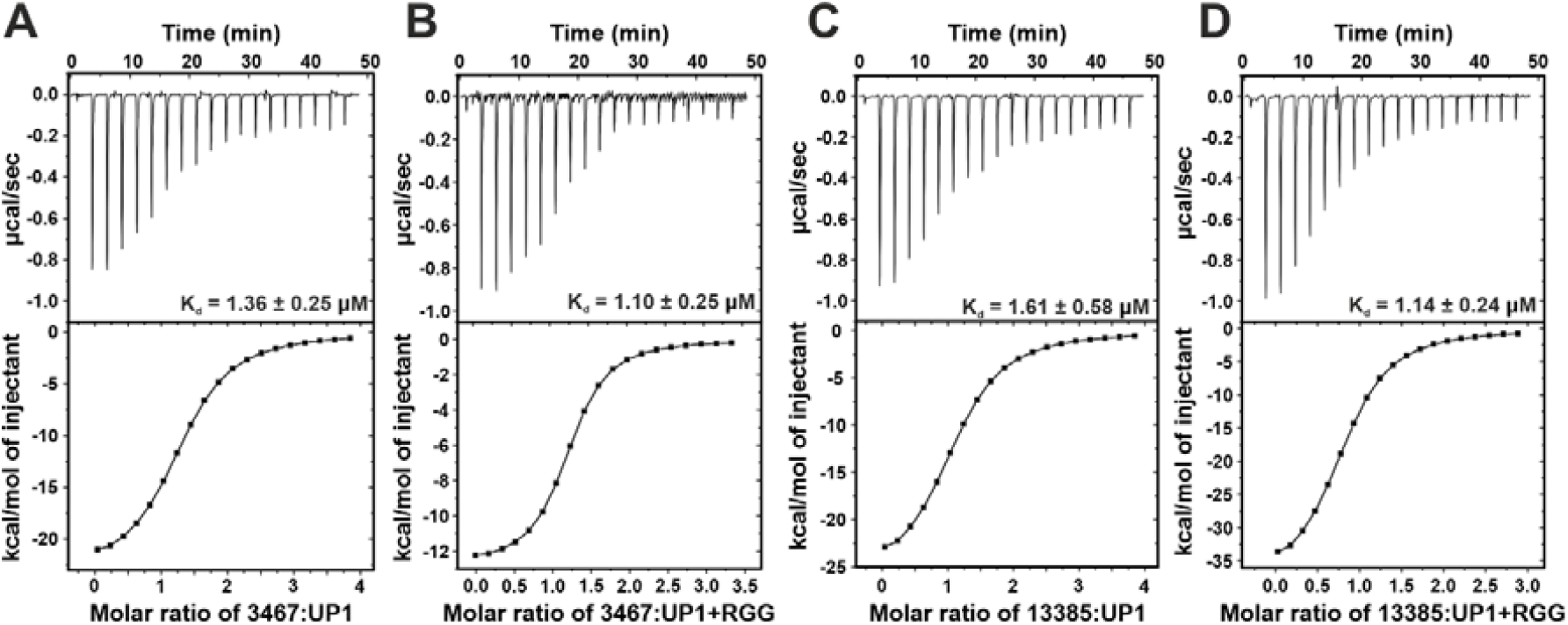
Thermodynamic characterisation of UP1 and UP1+RGG binding to the viral RNA G-quadruplexes using isothermal titration calorimetry. Raw thermograms (upper panels) and integrated binding isotherms fitted to a one-site binding model (lower panels) for titration of RNA in the sample cell with protein from the syringe are shown. **(A)** rG4-3467 titrated with UP1. **(B)** rG4-3467 titrated with UP1+RGG. **(C)** rG4-13385 titrated with UP1. **(D)** rG4-13385 titrated with UP1+RGG. The exothermic heat changes observed in all four titrations indicate enthalpically favourable binding. Dissociation constants (K_d_) obtained are shown with error from the fit on the corresponding thermogram.

Our ITC data demonstrate that the UP1 domain is the primary determinant for recognition of the SARS-CoV-2 RNA G-quadruplexes rG4-3467 and rG4-13385 by hnRNPA1. The comparable binding affinities of UP1 and UP1+RGG for both rG4s indicate that inclusion of the RGG-box does not significantly enhance binding under the conditions tested. This contrasts with its reported contribution to telomeric DNA and TERRA G4 recognition, suggesting that the role of the RGG-box may be G4-context dependent, whereas UP1 is sufficient for high-affinity recognition of these viral RNA G-quadruplexes.

### Remodelling of RNA G-quadruplex rG4-3467 and rG4-13385 by UP1 and UP1+RGG domains of hnRNPA1

In the next sets of experiments, we probed the possible remodelling and unfolding of the rG4-3467 and rG4-13385 SARS-CoV-2 G-quadruplexes by the UP1 and UP1+RGG domains using CD spectroscopy and ThT fluorescence experiments.

Increasing concentrations of UP1 and UP1+RGG progressively decreased the ellipticity at 264 nm of the rG4-3467 G-quadruplex, indicating unfolding of the G4 structure (Figure 4A and 4B). At an RNA:UP1 molar ratio of 1:2, we observed ∼32% decrease in the ellipticity at 264 nm of rG4-3467 G-quadruplex compared to RNA alone (without UP1 or UP1+RGG) (Figure 4A and 4B). Similarly, we observed unfolding of the rG4-13385 RNA G-quadruplex by the UP1 and UP1+RGG domains to a similar extent (Figure 4C and 4D).

**Figure 4.**
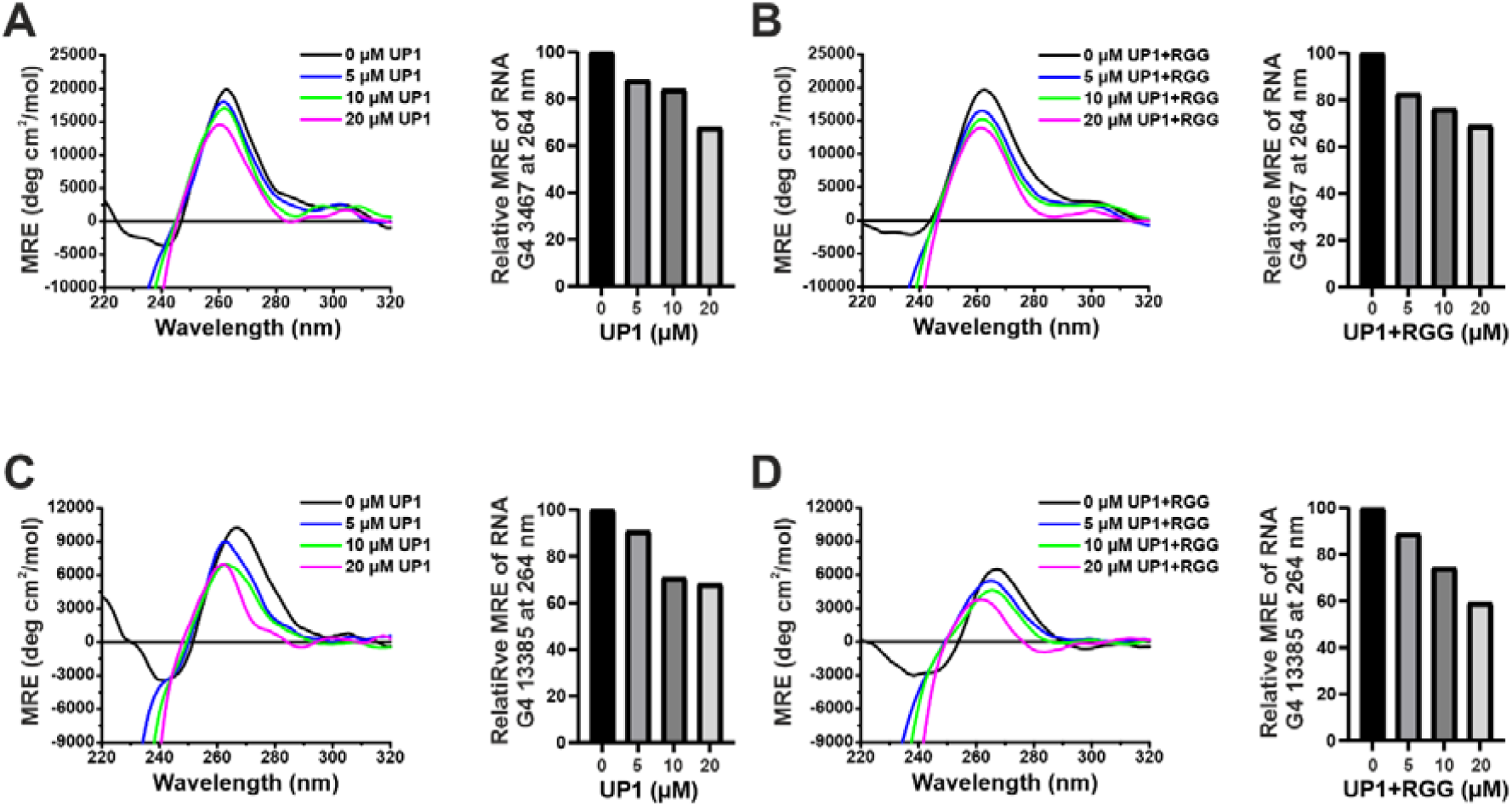
Titration of UP1 and UP1+RGG unfolds RNA G-quadruplex rG4-3467 and rG4-13385 as monitored by circular dichroism (CD). **(A, B)** CD spectra of rG4-3467 RNA in the presence of increasing concentrations 5 μM (blue), 10 μM (green) and 20 μM (magenta) UP1 **(A)** and UP1+RGG **(B). (C, D)** CD spectra of rG4-13385 RNA in the presence of increasing concentrations of 5 μM (blue), 10 μM (green) and 20 μM (magenta) UP1 **(C)** and UP1+RGG **(D)**. The corresponding bar graphs show the relative mean residue ellipticity (MRE) at 264 nm, normalized to the value obtained in the absence of protein.

The G-quadruplex unfolding was also monitored using the ThT fluorescence assay. Addition of UP1 or UP1+RGG to rG4-3467 (at 1:1 molar ratio) resulted in a reduction of the fluorescence intensity at ∼485 nm by ∼26% compared to RNA alone (without UP1 or UP1+RGG) (Figure 5A and 5B). In the case of rG4-13385 (at a 1:1 RNA:protein ratio), addition of UP1 and UP1+RGG resulted in an enhanced reduction of the fluorescence intensity at ∼485 nm by ∼37% and ∼50% respectively, compared to RNA alone (without UP1 or UP1+RGG) (Figure 5C and 5D).

**Figure 5.**
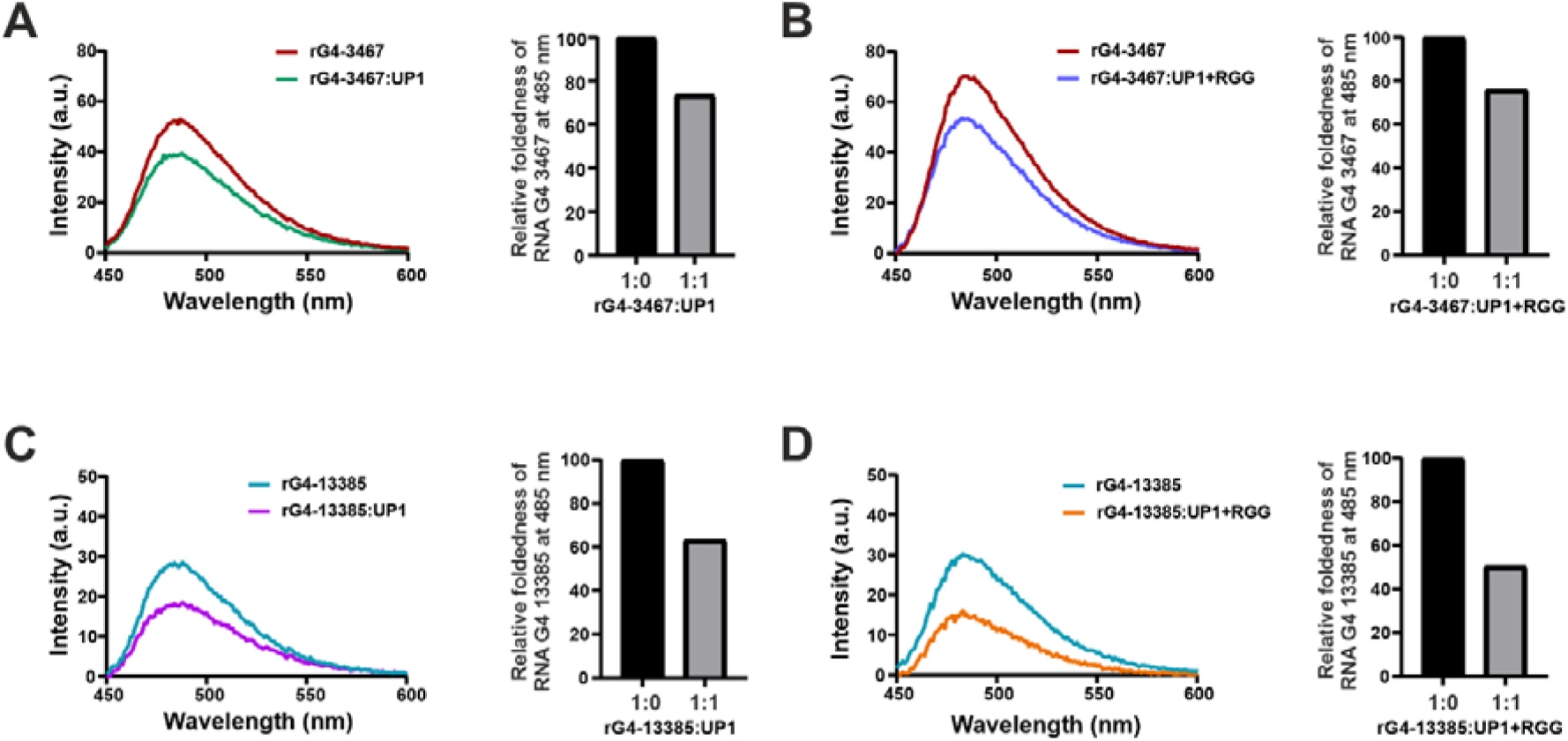
Unfolding of RNA G-quadruplexes rG4-3467 and rG4-13385 by UP1 and UP1+RGG monitored by Thioflavin T (ThT) fluorescence assay. Pre-folded RNA G-quadruplexes with ThT either alone (1:0) or with an equimolar concentration of protein (1:1), and steady-state emission was recorded from 450–600 nm following excitation at 425 nm. **(A, B)** Fluorescence emission spectra of rG4-3467 alone (red) and in the presence of UP1 (green) **(A)** and UP1+RGG (blue) **(B). (C, D)** Fluorescence emission spectra of rG4-13385 alone (cyan) and in the presence of UP1 (purple) **(C)** and UP1+RGG (orange) **(D)**. The corresponding bar graphs show the relative fold changes after addition of protein in fluorescence intensity at 485 nm, normalized to the RNA-only control (1:0). The RNA samples were prepared at a 1:1 molar ratio. Spectra are the average of three scans, baseline-corrected against buffer.

Taken together, the CD and ThT fluorescence results show that both UP1 and UP1+RGG destabilize the folded structures of rG4-3467 and rG4-13385, indicating that UP1 alone is sufficient to promote remodeling of these SARS-CoV-2 rG4s. Together with the comparable binding affinities observed by ITC, these findings support UP1 as the primary hnRNPA1 domain responsible for the recognition and structural remodeling of these viral RNA G-quadruplexes.

## Conclusions

Host RNA-binding proteins are increasingly recognised as determinants of the folding state of viral G-quadruplexes [24,28,33,34]. Here, we identify human hnRNPA1 as a binder and remodeller of the viral G-quadruplexes. We showed that hnRNPA1 binds and unfolds two SARS-CoV-2 rG4s located at the genomic positions 3467 and 13385. Interestingly, in previous studies, position 13385 was confirmed spectroscopically as an RNA G4 and shown to be bound by the viral helicase nsp13 [24], and position 3467, in the 5′ region of ORF1a, was shown to be stabilised by pyridostatin with consequent suppression of reporter translation [20].

These findings place hnRNPA1 within a set of proteins such as nsp13, nsp3, and CNBP which bind and remodel SARS-CoV-2 RNA G-quadruplexes leading to their stabilization or destabilization [24,28,33]. The folding state of a viral G-quadruplex is thus likely set by competition between stabilising and destabilising factors of both viral and host origin, as established at the viral G-quadruplexes such as HIV-1 LTR, where nucleolin and hnRNP A2/B1 exert opposing effects [60,61]. Therefore, we propose hnRNPA1 as a plausible regulator of the folding state of SARS-CoV-2 RNA G-quadruplexes in the host cell.

## Acknowledgments

The authors acknowledge funding for infrastructural support from the following programs of the Government of India: DST FIST, DST, DBT SAIF, UGC CAS, and the DBT-IISc partnership program.

Authors acknowledge funding support from the Anusandhan National Research Foundation (ANRF), Department of Science and Technology (DST), India (STAR award number STR/2021/000015 and CRG grant number CRG/2021/003597 to M.S.) and Ministry of Education (MoE). The research in the M.S. lab is supported by the grants from DBT (grant number BT/PR56485/BMS/85/598/2024), ICMR (file number EMDR/SG/10/2025-01384), and ANRF (ARG grant number ANRF/ARG/2025/007912/LS), India.

## Author contributions

M.S. conceived the research and acquired the funds. M.S. and P.G. designed the experiments. P.G. and S.M. performed all the experiments. A.M perform the bioinformatic analysis. The manuscript was written by all the authors.

## Data availability

This study did not generate unique reagents. All the materials of this study are available from the lead contact upon reasonable request.

## References

1 Varshney D, Spiegel J, Zyner K, Tannahill D & Balasubramanian S (2020) The regulation and functions of DNA and RNA G-quadruplexes. Nat Rev Mol Cell Biol 21, 459–474.

2 Phan AT, Kuryavyi V & Patel DJ (2006) DNA architecture: from G to Z. Curr Opin Struct Biol 16, 288–298.

3 Biffi G, Tannahill D, McCafferty J & Balasubramanian S (2013) Quantitative visualization of DNA G-quadruplex structures in human cells. Nat Chem 5, 182–186.

4 Biffi G, Di Antonio M, Tannahill D & Balasubramanian S (2014) Visualization and selective chemical targeting of RNA G-quadruplex structures in the cytoplasm of human cells. Nat Chem 6, 75–80.

5 Siddiqui-Jain A, Grand CL, Bearss DJ & Hurley LH (2002) Direct evidence for a G-quadruplex in a promoter region and its targeting with a small molecule to repress c-MYC transcription. Proc Natl Acad Sci 99, 11593–11598.

6 Kumari S, Bugaut A, Huppert JL & Balasubramanian S (2007) An RNA G-quadruplex in the 5′ UTR of the NRAS proto-oncogene modulates translation. Nat Chem Biol 3, 218– 221.

7 Cammas A, Dubrac A, Morel B, Lamaa A, Touriol C, Teulade-Fichou M-P, Prats H & Millevoi S (2015) Stabilization of the G-quadruplex at the VEGF IRES represses cap-independent translation. RNA Biol 12, 320–329.

8 Huppert JL & Balasubramanian S (2007) G-quadruplexes in promoters throughout the human genome. Nucleic Acids Res 35, 406–413.

9 Ghosh M & Singh M (2018) RGG-box in hnRNPA1 specifically recognizes the telomere G-quadruplex DNA and enhances the G-quadruplex unfolding ability of UP1 domain. Nucleic Acids Res 46, 10246–10261.

10 Beaudoin J-D & Perreault J-P (2010) 5’-UTR G-quadruplex structures acting as translational repressors. Nucleic Acids Res 38, 7022–7036.

11 Didiot M-C, Tian Z, Schaeffer C, Subramanian M, Mandel J-L & Moine H (2008) The G-quartet containing FMRP binding site in FMR1 mRNA is a potent exonic splicing enhancer. Nucleic Acids Res 36, 4902–4912.

12 Gomez D (2004) Telomerase downregulation induced by the G-quadruplex ligand 12459 in A549 cells is mediated by hTERT RNA alternative splicing. Nucleic Acids Res 32, 371–379.

13 Ruggiero E & Richter SN (2020) Viral G-quadruplexes: New frontiers in virus pathogenesis and antiviral therapy. In pp. 101–131.

14 Fleming AM, Ding Y, Alenko A & Burrows CJ (2016) Zika Virus Genomic RNA Possesses Conserved G-Quadruplexes Characteristic of the Flaviviridae Family. ACS Infect Dis 2, 674–681.

15 Métifiot M, Amrane S, Litvak S & Andreola M-L (2014) G-quadruplexes in viruses: function and potential therapeutic applications. Nucleic Acids Res 42, 12352–12366.

16 Wang S-R, Zhang Q-Y, Wang J-Q, Ge X-Y, Song Y-Y, Wang Y-F, Li X-D, Fu B-S, Xu G-H, Shu B, Gong P, Zhang B, Tian T & Zhou X (2016) Chemical Targeting of a G-Quadruplex RNA in the Ebola Virus L Gene. Cell Chem Biol 23, 1113–1122.

17 Qin G, Zhao C, Liu Y, Zhang C, Yang G, Yang J, Wang Z, Wang C, Tu C, Guo Z, Ren J & Qu X (2022) RNA G-quadruplex formed in SARS-CoV-2 used for COVID-19 treatment in animal models. Cell Discov 8, 86.

18 Lavezzo E, Berselli M, Frasson I, Perrone R, Palù G, Brazzale AR, Richter SN & Toppo S (2018) G-quadruplex forming sequences in the genome of all known human viruses: A comprehensive guide. PLOS Comput Biol 14, e1006675.

19 Ruggiero E & Richter SN (2018) G-quadruplexes and G-quadruplex ligands: targets and tools in antiviral therapy. Nucleic Acids Res 46, 3270–3283.

20 Razzaq M, Han JH, Ravichandran S, Kim J, Bae J-Y, Park M-S, Kannappan S, Chung W-C, Ahn J-H, Song MJ & Kim KK (2023) Stabilization of RNA G-quadruplexes in the SARS-CoV-2 genome inhibits viral infection via translational suppression. Arch Pharm Res 46, 598–615.

21 Zarudnaya M, Voiteshenko I, Hurmah V, Shyryna T, Nyporko A, Platonov M, Roszak S, Rasulev B, Kapusta K & Gorb L (2026) In Search of the Most Significant Potential G-Quadruplexes in SARS-CoV-2 RNA: Genomic Analysis. Viruses 18, 253.

22 Kabbara A, Vialet B, Marquevielle J, Bonnafous P, Mackereth CD & Amrane S (2022) RNA G-quadruplex forming regions from SARS-2, SARS-1 and MERS coronoviruses. Front Chem 10.

23 Cervenak M, Molnár OR, Horváth P & Smeller L (2024) Stabilization of G-Quadruplex Structures of the SARS-CoV-2 Genome by TMPyP4, BRACO19, and PhenDC3. Int J Mol Sci 25, 2482.

24 Ji D, Juhas M, Tsang CM, Kwok CK, L.Y & Zhang Y (2021) Discovery of G-quadruplex-forming sequences in SARS-CoV-2. Brief Bioinform 22, 1150–1160.

25 Nguyen LTA, Nguyen TTT & Dang DT (2024) Specific binding of G-quadruplex in SARS-CoV-2 RNA by RHAU peptide. Curr Res Struct Biol 7, 100126.

26 Cui H & Zhang L (2020) G-Quadruplexes Are Present in Human Coronaviruses Including SARS-CoV-2. Front Microbiol 11.

27 Zhang R, Xiao K, Gu Y, Liu H & Sun X (2020) Whole Genome Identification of Potential G-Quadruplexes and Analysis of the G-Quadruplex Binding Domain for SARS-CoV-2. Front Genet 11.

28 Bezzi G, Piga EJ, Binolfi A & Armas P (2021) CNBP Binds and Unfolds In Vitro G-Quadruplexes Formed in the SARS-CoV-2 Positive and Negative Genome Strands. Int J Mol Sci 22, 2614.

29 Panera N, Tozzi AE & Alisi A (2020) The G-Quadruplex/Helicase World as a Potential Antiviral Approach Against COVID-19. Drugs 80, 941–946.

30 Belmonte-Reche E, Serrano-Chacón I, Gonzalez C, Gallo J & Bañobre-López M (2021) Potential G-quadruplexes and i-Motifs in the SARS-CoV-2. PLoS One 16, e0250654.

31 Zhao C, Qin G, Niu J, Wang Z, Wang C, Ren J & Qu X (2021) Targeting RNA G-Quadruplex in SARS-CoV-2: A Promising Therapeutic Target for COVID-19? Angew Chemie Int Ed 60, 432–438.

32 Liu G, Du W, Sang X, Tong Q, Wang Y, Chen G, Yuan Y, Jiang L, Cheng W, Liu D, Tian Y & Fu X (2022) RNA G-quadruplex in TMPRSS2 reduces SARS-CoV-2 infection. Nat Commun 13, 1444.

33 Lavigne M, Helynck O, Rigolet P, Boudria-Souilah R, Nowakowski M, Baron B, Brülé S, Hoos S, Raynal B, Guittat L, Beauvineau C, Petres S, Granzhan A, Guillon J, Pratviel G, Teulade-Fichou M-P, England P, Mergny J-L & Munier-Lehmann H (2021) SARS-CoV-2 Nsp3 unique domain SUD interacts with guanine quadruplexes and G4-ligands inhibit this interaction. Nucleic Acids Res 49, 7695–7712.

34 Burghaus M, Staschko T, Mendes M, Filcenkova L, Gerhard J, Dittmar G, Ihmels H & Krauß S (2026) A G-quadruplex structure in the SARS-CoV-2 RNA recruits human topoisomerase I. Front RNA Res 3.

35 Jean-Philippe J, Paz S & Caputi M (2013) hnRNP A1: The Swiss Army Knife of Gene Expression. Int J Mol Sci 14, 18999–19024.

36 Roy R, Huang Y, Seckl MJ & Pardo OE (2017) Emerging roles of <scp>hnRNPA1</scp> in modulating malignant transformation. WIREs RNA 8.

37 Hudson JS, Ding L, Le V, Lewis E & Graves D (2014) Recognition and Binding of Human Telomeric G ⍰ Quadruplex DNA by Unfolding Protein 1.

38 Pollard VW, Michael WM, Nakielny S, Siomi MC, Wang F & Dreyfuss G (1996) A Novel Receptor-Mediated Nuclear Protein Import Pathway. Cell 86, 985–994.

39 Nishikawa T, Kuwano Y, Takahara Y, Nishida K & Rokutan K (2019) HnRNPA1 interacts with G-quadruplex in the TRA2B promoter and stimulates its transcription in human colon cancer cells. Sci Rep 9, 10276.

40 Ma Y, Yang Y, Xin J, He L, Hu Z, Gao T, Pan F & Guo Z (2023) RNA G-Quadruplex within the 5′-UTR of FEN1 Regulates mRNA Stability under Oxidative Stress. Antioxidants 12, 276.

41 Michlewski G, Guil S, Semple CA & Cáceres JF (2008) Posttranscriptional Regulation of miRNAs Harboring Conserved Terminal Loops. Mol Cell 32, 383–393.

42 Guil S & Cáceres JF (2007) The multifunctional RNA-binding protein hnRNP A1 is required for processing of miR-18a. Nat Struct Mol Biol 14, 591–596.

43 Cammas A, Lacroix-Triki M, Pierredon S, Le Bras M, Iacovoni JS, Teulade-Fichou M-P, Favre G, Roché H, Filleron T, Millevoi S & Vagner S (2016) hnRNP A1-mediated translational regulation of the G quadruplex-containing RON receptor tyrosine kinase mRNA linked to tumor progression. Oncotarget 7, 16793–16805.

44 Cortolezzis Y, Othman Z, Agostini F, Ibrahim I, Picco R, Salgado GF, D. Giorgio E & Xodo LE (2025) Post-transcriptional control of KRAS?: functional roles of 5′UTR RNA G-quadruplexes, long noncoding RNA, and hnRNPA1. Nucleic Acids Res 53.

45 Ghosh M & Singh M (2020) Structure specific recognition of telomeric repeats containing RNA by the RGG-box of hnRNPA1. Nucleic Acids Res 48, 4492–4506.

46 Paramasivam M, Membrino A, Cogoi S, Fukuda H, Nakagama H & Xodo LE (2009) Protein hnRNP A1 and its derivative Up1 unfold quadruplex DNA in the human KRAS promoter: implications for transcription. Nucleic Acids Res 37, 2841–2853.

47 Dabral P, Babu J, Zareie A & Verma SC (2020) LANA and hnRNP A1 Regulate the Translation of LANA mRNA through G-Quadruplexes. J Virol 94.

48 Jing H, Liu Y, Peng Z, Duan E, Liu J, Lv Y, Sun Y, Dong W, Li X, Wang J, Cao S, Wang H, Zhang Y & Li H (2025) The UP1 domain is essential for the facilitation effect of HnRNP A1 on PRRSV-2 replication. Virology 603, 110378.

49 Huang P & Lai MMC (2001) Heterogeneous Nuclear Ribonucleoprotein A1 Binds to the 3′-Untranslated Region and Mediates Potential 5′-3′-End Cross Talks of Mouse Hepatitis Virus RNA. J Virol 75, 5009–5017.

50 Li H-P, Zhang X, Duncan R, Comai L & Lai MMC (1997) Heterogeneous nuclear ribonucleoprotein A1 binds to the transcription-regulatory region of mouse hepatitis virus RNA. Proc Natl Acad Sci 94, 9544–9549.

51 Schmidt N, Lareau CA, Keshishian H, Ganskih S, Schneider C, Hennig T, Melanson R, Werner S, Wei Y, Zimmer M, Ade J, Kirschner L, Zielinski S, Dölken L, Lander ES, Caliskan N, Fischer U, Vogel J, Carr SA, Bodem J & Munschauer M (2020) The SARS-CoV-2 RNA–protein interactome in infected human cells. Nat Microbiol 6, 339–353.

52 Kumar R, Khandelwal N, Chander Y, Nagori H, Verma A, Barua A, Godara B, Pal Y, Gulati BR, Tripathi BN, Barua S & Kumar N (2022) S-adenosylmethionine-dependent methyltransferase inhibitor DZNep blocks transcription and translation of SARS-CoV-2 genome with a low tendency to select for drug-resistant viral variants. Antiviral Res 197, 105232.

53 Sievers F, Wilm A, Dineen D, Gibson TJ, Karplus K, Li W, Lopez R, McWilliam H, Remmert M, Söding J, Thompson JD & Higgins DG (2011) Fast, scalable generation of high-quality protein multiple sequence alignments using Clustal Omega. Mol Syst Biol 7, MSB201175.

54 Kikin O, D’Antonio L & Bagga PS (2006) QGRS Mapper: a web-based server for predicting G-quadruplexes in nucleotide sequences. Nucleic Acids Res 34, W676– W682.

55 del Villar-Guerra R, Trent JO & Chaires JB (2018) G-Quadruplex Secondary Structure Obtained from Circular Dichroism Spectroscopy. Angew Chemie Int Ed 57, 7171–7175.

56 Renaud de la Faverie A, Guédin A, Bedrat A, Yatsunyk LA & Mergny J-L (2014) Thioflavin T as a fluorescence light-up probe for G4 formation. Nucleic Acids Res 42, e65–e65.

57 Sjekloca L & Ferré-D’Amaré AR (2019) Binding between G Quadruplexes at the Homodimer Interface of the Corn RNA Aptamer Strongly Activates Thioflavin T Fluorescence. Cell Chem Biol 26, 1159-1168.e4.

58 Campbell NH, Parkinson GN, Reszka AP & Neidle S (2008) Structural Basis of DNA Quadruplex Recognition by an Acridine Drug. J Am Chem Soc 130, 6722–6724.

59 Parkinson GN, Ghosh R & Neidle S (2007) Structural Basis for Binding of Porphyrin to Human Telomeres,. Biochemistry 46, 2390–2397.

60 Tosoni E, Frasson I, Scalabrin M, Perrone R, Butovskaya E, Nadai M, Palù G, Fabris D & Richter SN (2015) Nucleolin stabilizes G-quadruplex structures folded by the LTR promoter and silences HIV-1 viral transcription. Nucleic Acids Res 43, 8884–8897.

61 Scalabrin M, Frasson I, Ruggiero E, Perrone R, Tosoni E, Lago S, Tassinari M, Palù G & Richter SN (2017) The cellular protein hnRNP A2/B1 enhances HIV-1 transcription by unfolding LTR promoter G-quadruplexes. Sci Rep 7, 45244.

