## Supplementary Information for "hnRNPA1 Remodels SARS-CoV-2 Genomic G-Quadruplexes through Its UP1 Domain"

**Supplementary Figure 1**

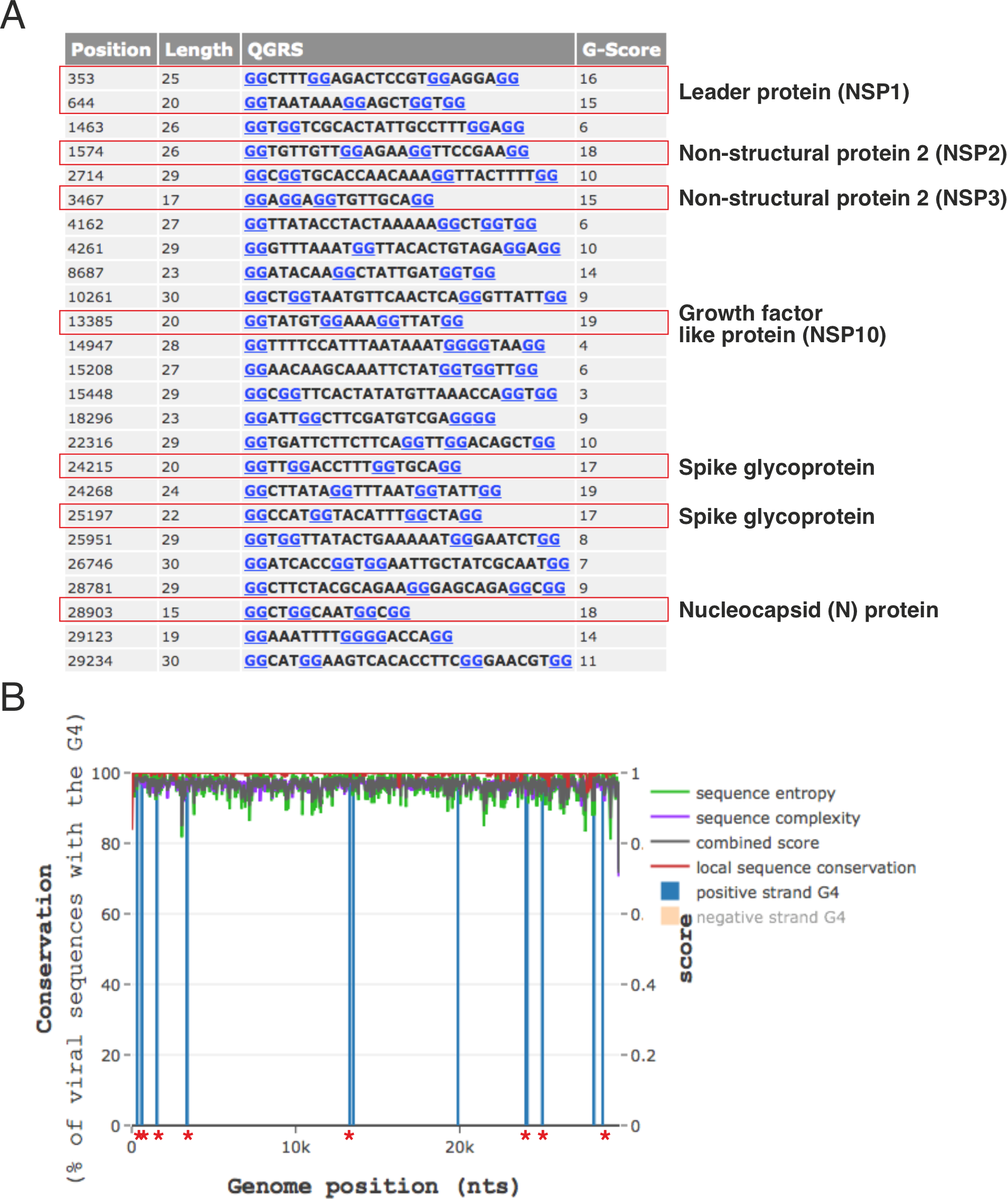

**Figure S1. Predicted putative G-quadruplex-forming sequences in the SARS-CoV-2 genome.** Output of QGRS Mapper for the SARS-CoV-2 Wuhan reference genome, showing the twenty-five non-overlapping putative quadruplex-forming sequences (PQSs). For each PQS the genomic position of the first nucleotide, the length of the predicted quadruplex-forming region, the sequence, and the G-score assigned by QGRS Mapper are listed; guanine residues predicted to participate in G-tetrad formation are shown in blue. The eight sequences found to be conserved among SARS-CoV-2 and related SARS coronaviruses are boxed in red, and the viral protein encoded at each of these positions is indicated on the right.

**Supplementary Figure 2**

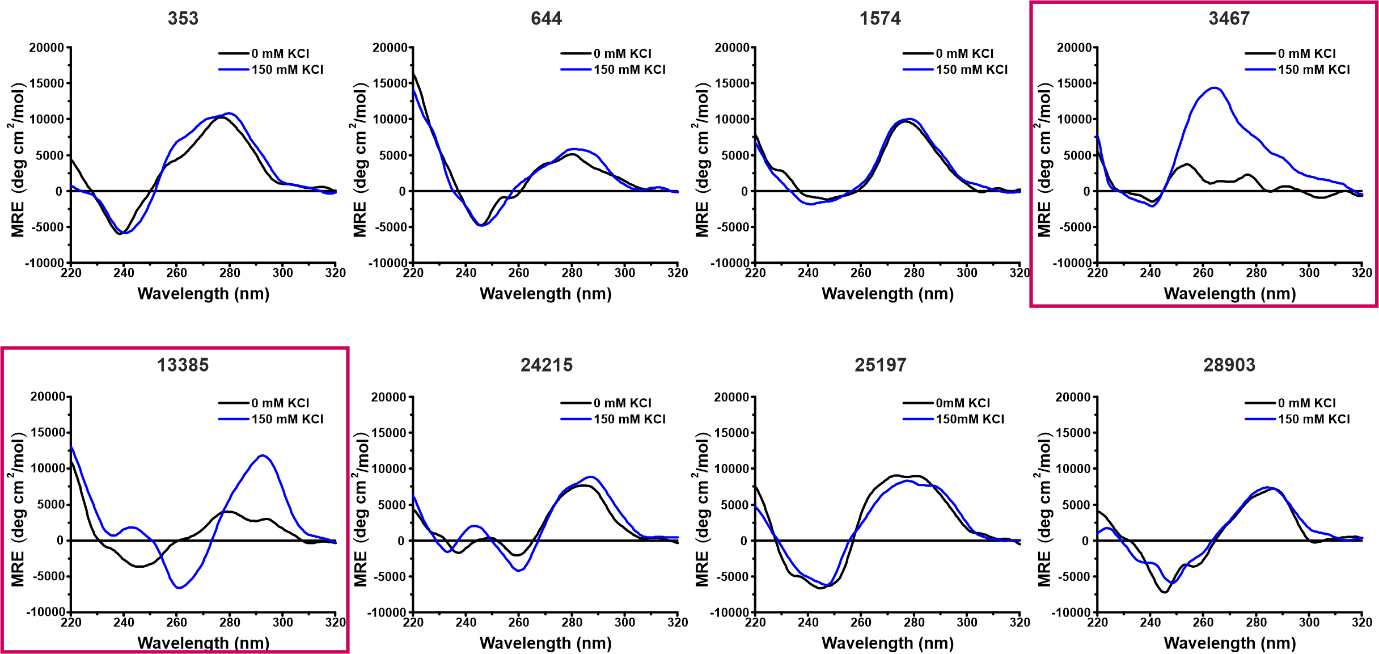

**Figure S2. K⁺ dependence of G-quadruplex formation by the eight conserved DNA oligonucleotides.** CD spectra of each of the eight DNA oligonucleotides recorded from 220 to 320 nm in the absence (0 mM, black) and presence (100 mM, blue) of KCl. Sequence identifiers are indicated above each panel. Only dG4-3467 and dG4-13385, boxed in red, show a significant K⁺-dependent change in their spectra: in the presence of KCl, dG4-3467 develops a positive maximum at ~264 nm characteristic of a parallel G-quadruplex, and dG4-13385 develops a positive maximum at ~295 nm with a negative minimum at ~264 nm characteristic of an antiparallel G-quadruplex, neither of which is present at 0 mM KCl. The remaining six sequences show superimposable spectra at both salt concentrations, indicating that their conformations are not K⁺-dependent and that they do not form g-quadruplex structures under these conditions.

**Supplementary Figure 3**

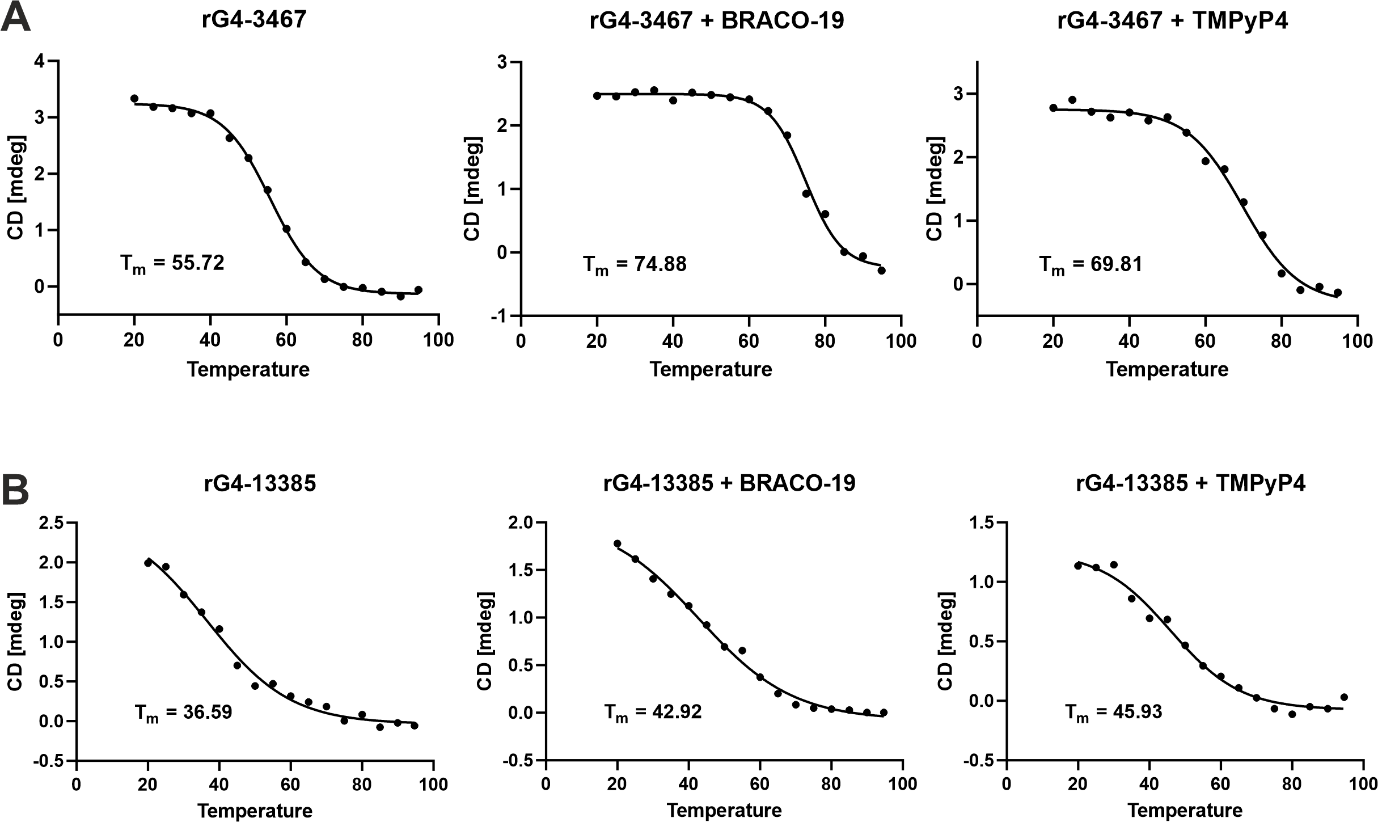

**Figure S3. Thermal stabilisation of the viral RNA G-quadruplexes by BRACO-19 and TMPyP4.** Thermal melting profiles of rG4-3467 (A) and rG4-13385 (B) monitored by CD, alone and in the presence of an equimolar concentration of the G-quadruplex-stabilising ligands BRACO-19 and TMPyP4. Ellipticity at 264 nm was recorded from 20 to 95 °C in 5 °C intervals and melting temperatures (T_m_) were obtained by fitting the melting curves to a Boltzmann sigmoidal equation; fitted curves are shown as solid lines and the derived T_m_ is indicated on each panel. **(A)** For rG4-3467, T_m_ increases from 55.7 °C to 74.9 °C with BRACO-19 and to 69.8 °C with TMPyP4. **(B)** For rG4-13385, T_m_ increases from 36.6 °C to 42.9 °C with BRACO-19 and to 45.9 °C with TMPyP4.
